# SNP-guided identification of monoallelic DNA-methylation events from enrichment-based sequencing data

**DOI:** 10.1101/002352

**Authors:** Sandra Steyaert, Wim Van Criekinge, Ayla De Paepe, Simon Denil, Klaas Mensaert, Katrien Vandepitte, Wim Vanden Berghe, Geert Trooskens, Tim De Meyer

**Affiliations:** Department of Mathematical Modelling, Statistics and Bioinformatics, University of Ghent, Belgium; Department of Biology, University of Leuven, Belgium; PPES, Department of Biomedical Sciences, University of Antwerp, Belgium

**Keywords:** Monoallelic events, DNA-methylation, SNPs, enrichment-based sequencing, imprinting

## Abstract

Monoallelic gene expression is typically initiated early in the development of an organism. Dysregulation of monoallelic gene expression has already been linked to several non-Mendelian inherited genetic disorders. In humans, DNA-methylation is deemed to be an important regulator of monoallelic gene expression, but only few examples are known. One important reason is that current, cost-affordable truly genome-wide methods to assess DNA-methylation are based on sequencing post enrichment. Here, we present a new methodology that combines methylomic data from MethylCap-seq with associated SNP profiles to identify monoallelically methylated loci. Using the Hardy-Weinberg theorem for each SNP locus, it could be established whether the observed frequency of samples featured by biallelic methylation was lower than randomly expected. Applied on 334 MethylCap-seq samples of very diverse origin, this resulted in the identification of 80 genomic regions featured by monoallelic DNA-methylation. Of these 80 loci, 49 are located in genic regions of which 25 have already been linked to imprinting. Further analysis revealed statistically significant enrichment of these loci in promoter regions, further establishing the relevance and usefulness of the method. Additional validation of the found loci was done using 14 whole-genome bisulfite sequencing data sets. Importantly, the developed approach can be easily applied to other enrichment-based sequencing technologies, such as the ChIP-seq-based identification of monoallelic histone modifications.

## INTRODUCTION

For diploid organisms, gene expression is denoted as monoallelic if only one allele is transcriptionally active. The expressed allele can be randomly selected (e.g. X-chromosome inactivation and some autosomal genes) or predetermined by parental imprinting (Gimelbrant et al. 2007; Ferguson-Smith 2011; Fedoriw et al. 2012). Erroneous monoallelic expression has been associated to several genetic disorders, like the Prader-Willi syndrome, as well as to certain forms of cancer, like Wilms’ tumor. Both diseases are caused by loss of imprinting of some genes in the 15q11-q13 and 11p15.5 region, respectively (Egger et al. 2004). Epigenetics is defined as the study of inheritable modifications on both chromatin and DNA that have an influence on gene expression without changing the underlying DNA sequence (Goldberg et al. 2007). Mammalian DNA-methylation is an epigenetic mark that is predominantly found in a CpG sequence context (Law et al. 2010). This methylation mark has been linked with gene expression and when located in the promoter region, it generally leads to transcriptional silencing of the corresponding gene (Jones 2012). As it is a defining feature of cellular identity and essential for normal development, its dysregulation is often associated with disease (Egger et al. 2004). Monoallelic DNA-methylation is likely to bare an important role in the regulation of monoallelic expression (Milani et al. 2009). In addition to DNA-methylation, histone modifications also contribute to the maintenance of monoallelic expression. The methylated, silenced allele is mostly sustained with the repressive histone modification histone H3 trimethylation at lysine 9 (H3K9me3) while the active allele is characterized by the permissive histone marker H3 trimethylation at lysine 4 (H3K4me3) (Kacem and Feil 2009).

Although imprinting is a well-investigated topic and several studies already provided evidence (e.g. computational predictions based on DNA sequence characteristics or detection of monoallelic expression) of some regions with monoallelic DNA-methylation (Luedi et al. 2005, Luedi et al. 2007, Babak et al. 2008, Serre et al. 2008, Wang et al. 2008, Morison et al. 2001; Zhang et al. 2010; Ferguson-Smith 2011; Fedoriw et al. 2012), only a few imprinted regions are well characterized in humans, like for example the *IGF2/H19* region. Furthermore, while monoallelic methylation has been shown to play an important role in the differentiation between tissues, little is known about the presence of loci that are stably monoallelically methylated in all tissues as well as the genome-wide character of monoallelic DNA-methylation. The recent advent of next-generation massively parallel sequencing platforms has introduced the possibility of genome-wide DNA-methylation profiling. Bisulfite sequencing, which combines bisulfite treatment of genomic DNA with the high-throughput sequencing of the entire genome, is the gold standard and allows to readily identify monoallelic methylated alleles (Shoemaker et al. 2010), but is very costly and therefore outside the reach of smaller projects. Fortunately, cost-effective alternatives based on the specific enrichment of methylated portions of the genome (i.e. enrichment-based methods) such as methylated DNA immunoprecipitation followed by sequencing (MeDIP-seq) and methyl-CpG binding domain protein sequencing (MethylCap-seq) exist. Yet, these methods do neither provide single base pair resolution nor information regarding unmethylated alleles and are therefore not directly applicable to detect monoallelic events (Serre et al. 2010). While some approaches already tried to tackle this issue, they rely on the combination of multiple sequencing technologies, like for example the integrative method of Harris et al. (2010), which tries to find regions with intermediate and potentially monoallelic events by combining data originating from MeDIP-seq, methylation-sensitive restriction enzyme sequencing (MRE-seq), ribonucleic acid sequencing (RNA-seq) and chromatin immunoprecipitation followed by sequencing (ChIP-seq).

To circumvent these issues, we developed a data-analytical framework that solely uses data from enrichment-based sequencing (like MethylCap-seq), which screens for regions that exhibit monoallelic DNA-methylation based on classical population genetic theory. The developed pipeline first compares enrichment-based sequencing data of multiple samples to the public NCBI Single Nucleotide Polymorphism (SNP)-archive (dbSNP) in order to screen the obtained non-duplicate, uniquely mappable sequence reads for SNPs. Only SNP loci with an adequately coverage and allele frequency are retained and the effect of sequencing errors is further reduced by comparing the chance of a sequencing error with the chance of detecting genuine SNPs. For each single SNP locus, the Hardy-Weinberg theorem is then applied to evaluate whether the observed frequency of samples featured by a biallelic event is lower than randomly expected (Mayo 2008). Using a permutation approach, confidence limits are simulated and genomic regions with a p-value smaller than the p-value corresponding with a given false discovery rate (FDR) can be assumed to harbour a monoallelic event.

Starting from MethylCap-seq data of a mixture of 334 Caucasian human samples and an FDR of 0.1, this methodology allowed the identification of 80 monoallelically methylated loci, for which significant enrichment was found in promoter regions. Of these 80 loci, 25 have previously been linked to imprinting. Additional validation of 44 of the found loci was done using 14 whole-genome bisulfite sequencing (WGBS) data sets. 29 loci (65.9%) showed evidence of monoallelic methylation in at least one of these samples. This outcome as well as the developed approach are innovative and provide new insights. Because a variety of tissues was used in the analysis, the detected loci are likely generally imprinted in a large set of tissues. Furthermore, our method allows the identification of monoallelically methylated loci in a parental-independent and genome-wide manner. Finally, because of the general rationale of the developed approach, it can be applied to enrichment-based sequencing applications to detect monoallelic features other than DNA-methylation. A possible application could be ChIP-seq (Furey 2012) to screen for monoallelic histone modifications (Kadota et al. 2007, Maynard et al. 2008, Birney et al. 2010, Kasowski et al. 2010, McDaniell et al. 2010).

## METHODS

### 1. Samples

A total of 334 human samples, mostly cancer samples of various tissues, was used to detect monoallelically methylated loci (Supplemental Table 1.1). Of these 334 samples, 215 samples were of female origin and only these were used to analyse the X-chromosome. Genomic DNA (gDNA) was extracted from these samples with the Easy DNA kit (Invitrogen K1800-01) using protocol #4 from the manufacturer manual. The DNA-concentration was measured on a Nanodrop ND-1000. Subsequently, the gDNA was sheared on the Covaris S2 with following settings: duty cycle 10%, intensity 5, 200 cycles per burst during 180 seconds to obtain fragments with an average length of 200 bp. The power mode was frequency sweeping, temperature 6-8°C and water level of 12. 500 ng was loaded in 130 μl TE (1:5) in a microtube with AFA intensifier.

### 2. Methyl-CpG binding domain sequencing

Methyl-CpG binding domain protein sequencing (MethylCap-seq) (Serre et al. 2010), which combines enrichment of methylated DNA-fragments by MBD-based affinity purification with massively parallel sequencing, was used to profile the DNA-methylation pattern of the 334 samples. The samples were sequenced according to the protocol described in the paper of De Meyer et al. (2013) with some additional modifications: i) After DNA-fragmentation, the methylated fragments were captured using Diagenode’s MethylCap kit starting from a DNA-concentration of 500 ng instead of 200 ng. ii) Paired-end sequencing was done on either the Illumina GAIIx or the HiSeq platform. Depending on the sequencing platform, the obtained paired-end sequence reads were 45 or 51 bp, respectively.

### 3. DATA PRE-PROCESSING

The rationale behind the proposed methodology is that biallelic DNA-methylation results in MethylCap-seq data which is in Hardy-Weinberg equilibrium for each locus, i.e. if SNPs are present for a locus, both homozygous and heterozygous subjects will be detected at a predictable rate (Mayo 2008). However, in case of monoallelic methylation, heterozygous samples will no longer be detected resulting in deviation from the Hardy-Weinberg equilibrium, which can be measured. For a detailed description of the statistical framework, see the Supplemental Methods. Figure 1 gives an overall representation of the workflow starting from MethylCap-seq data.

**Figure 1.**
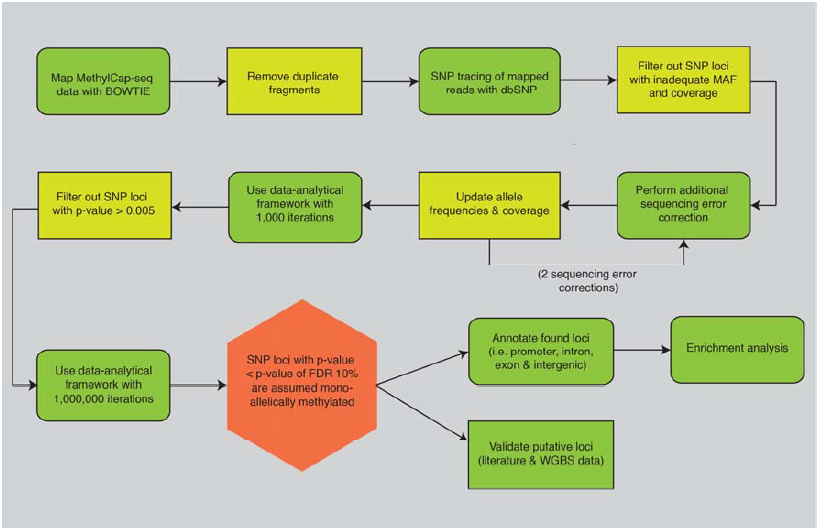
Overview of the developed bioinformatics pipeline to detect putative monoallelically methylated SNP loci starting from MethylCap-seq data. After mapping with BOWTIE the non-duplicate, uniquely mapped reads are screened for SNPs using dbSNP. To reduce the computational load SNP loci with a too high major allele frequency (MAF) and/or a too low overall coverage are filtered. In this reduced data set, an additional sequencing error correction was performed with two iterations. The corrected data was next put in the newly developed data-analytical framework with 1,000 and 1,000,000 iterations, respectively. Only loci that obtained a p-value smaller than or equal to 0.005 after the first iteration were kept as input for the second iteration. If the p-value obtained for a locus was smaller than the p-value corresponding with an FDR of 0.1 the monoallelic methylation on this locus was called significant. After determining the functional annotation of these SNP positions an enrichment analysis was performed. Finally, the resulting loci were validated using both literature and whole-genome bisulfite sequencing (WGBS) data.

#### 3.1 Mapping

For each of the 334 samples, the MethylCap-seq paired-end reads were mapped using BOWTIE (Langmead et al. 2009). The mapping parameters were chosen so that only those paired-end reads that mapped uniquely on the human hg19/GRCh37 reference assembly within a maximum of 400 bp of each other were retained. In order to both reduce the presence of sequencing errors as well as to allow the occurrence of real SNPs, a maximum of only three mismatches was allowed. Duplicate fragments, i.e. fragments with the exact same location of both paired-end reads, were disposed as these are most likely the result of amplification of the same sequence reads during the library preparation.

#### 3.2 SNP tracing

The non-duplicate, uniquely mappable reads were subsequently screened for SNPs. Only positions that showed a mismatch in the mapping of one or more samples and that overlapped known single nucleotide variations (SNVs) of the Single Nucleotide Polymorphism Database of NCBI (dbSNP, version 137) were withheld. Not keeping all the mismatches reduces both the effect of sequencing errors (false positives) and the computational load in the further analyses. Also, for each locus, the coverage of each SNP variant was determined, and the allele frequencies were estimated.

#### 3.3 Additional data filtering and correction

Both for computational reasons and as a first filtering step for sequencing errors, SNP loci with a very high major allele frequency were filtered (threshold 0.9). Additionally, a minimal total coverage threshold, i.e. across all samples, for each SNP locus was imposed (350 ∼ 1x per sample). Note that loci not fulfilling both criteria are unlikely to provide sufficient power for the subsequent statistical analysis. As analysis of the X-chromosome involved fewer samples, the threshold for the coverage was set to a less stringent value, namely 250 instead of 350, which roughly corresponded to the number of female subjects.

In this reduced data set, an additional sequencing error correction was performed. For computational reasons, a simple Bayesian methodology was implemented. Basically, for each sample and locus, the chance of obtaining a certain profile was calculated under i) the assumption of heterozygosity and, ii) the assumption of homozygosity but with additional sequencing errors. The option with the largest a posteriori change was withheld (with alleles representing putative sequencing errors being removed from the data set). As the prior chances of homozygosity and heterozygosity were based on the allele frequencies, which are updated upon each round of the Bayesian algorithm, this method was performed twice (see Supplemental Methods section 2.1.3). This approach can be considered to be conservative (i.e. to disfavour the presence of monoallelic DNA-methylation), as i) only two rounds of correction were applied and, ii) the sequencing error estimate (0.25%, based on Quail et al. 2012) is on the lower bound of estimates reported and is based on the performance of the Illumina HiSeq, whereas also more error prone GAIIx data were included in this study.

### 4. Detection of monoallelically methylated loci

After additional filtering and data correction, the remaining data were used as input of the new data-analytical framework developed in the R statistical environment (R 2.15.2). The statistical strategy and practical implementation are elaborated in the Supplemental Methods (section 2.1). In summary, based on the observed allele frequencies, theoretically expected genotype frequencies can be calculated assuming Hardy-Weinberg equilibrium in the overall data set. If the observed frequency of heterozygote individuals is significantly reduced relative to Hardy-Weinberg expectations, this indicates significant monoallelic methylation. Null distributions were generated using random data with the same allele frequencies and sample coverages (for that locus) as in the original data. This approach accounts for the increased likelihood of erroneously calling loci with a low coverage homozygous. P-values were determined by comparison of the observed frequency of heterozygotes with the generated null distributions. Only loci that obtained a p-value smaller than or equal to 0.005 after the first iteration were kept as input for the second iteration. Thus, after the first iteration round, loci that were in all probability not monoallelically methylated, were filtered out as to reduce the computational time in the second iteration. At the end of the second iteration the algorithm obtained a p-value for each locus. If this p-value was smaller than the p-value corresponding with an FDR of 0.1, monoallelic methylation on this locus was called significant. This procedure was also performed two times, a first time with 1,000 and a second time with 1,000,000 iterations. To summarize results, significant loci were visualised on a circular plot with the Circos tool (Krzywinski et al. 2009).

### 5. Functional annotation and enrichment analysis

Successful completion of the monoallelically methylated loci detection pipeline resulted in a list of significant SNPs. The functional annotation (i.e. promoter, exon, intron and intergenic) of these SNP positions was determined using Ensembl (release 66), wherein the promoter was defined as starting from 2000 bp upstream until the transcriptional start site.

We tested for enrichment in one or more of these functional categories. A null distribution was generated by random sampling from the total amount of detected SNPs (after filtering as specified in Methods section 3.3) and counting the occurrences of the respective annotations. During this sampling procedure, the number of SNPs sampled for each chromosome was equal to the number of significant SNPs on that chromosome. This sampling was repeated 1,000 times. With the null distribution obtained for each of these functional locations (i.e. promoter, exon, intron and intergenic), it was possible to calculate a two-sided p-value for each functional location. For loci that were featured by more than one functional annotation (i.e. overlapping genes and/or different transcripts and/or sense and antisense strand) the score for the functional location was divided by the amount of different functional locations that this locus has (the sum always being one). For example, if a locus is located in an exon on the sense strand but is also located in an intron on the other strand, both the exon and intron were attributed a score of 0.5.

### 6. Validation of putative loci using 14 whole-genome bisulfite sequencing data sets

In order to evaluate the loci detected by this novel methodology, an extra validation step was performed using 14 publicly available WGBS data sets comprising a range of tissue types. The WGBS data sets were downloaded from the Gene Expression Omnibus (GEO) repository (Edgar et al. 2002). A summary of the data sets including accession numbers is provided in Supplemental Table 1.3. The 14 samples were aligned in a window of 2000 bp (1000 bp upstream and 1000 bp downstream) around the candidate SNP positions (hg19/GRCh37 reference assembly) using BISMARK (Krueger and Andrews 2011). After excluding duplicates, only reads mapping onto one of the candidate monoallelically SNP positions were kept. Next, for each SNP position and each sample the methylation calls of all CpGs were summarized per SNP allele (covered by the reads on the specific SNP position). To assess monoallelic DNA-methylation in the SNP loci a Pearson chi-square test was performed. Samples that were not covered or were homozygous for the particular locus were excluded. In summary, for each heterozygous sample a chi-square value was calculated based on the degree of (non-)methylation obtained for each SNP allele, with a high chi-square value indicating that the methylation degree is allele dependent (i.e. monoallelic methylation). Null distributions were made by a permutation approach (using the chisq.test function of the R Stats package) generating 2000 random chi-square values for each sample, making it possible to determine a sample-specific p-value for each SNP-loci. By summing the chi-square values over all heterozygous samples for a specific SNP locus and again generating null distributions of random chi-square values, also a global p-value for a SNP locus could be obtained. Note that this test does not require absolute absence of methylation of one allele, which would be too strict given the possibility of incomplete bisulfite conversion and the presence of both sequencing errors and sequencing bias.

## RESULTS

### Mapping

For the 334 samples the mapping resulted in 2,995,375,490 uniquely mapped reads and an average mapping percentage of 63.05% (Supplemental Table 1.1). After removing the duplicate fragments a total of 2,688,409,588 non-duplicate, uniquely mapped reads was acquired.

### SNP tracing and data filtering

After parsing the mapping output for SNPs (= mapping mismatches), 19,850,891 SNPs overlapped with already known SNV positions from dbSNP. These 19,850,891 loci represent 41.61% of the total number of SNV present in dbSNP and only these SNPs were used in the remainder of the analysis. Supplemental Table 1.2 details the number of SNPs that overlapped with dbSNP per chromosome.

After preprocessing the data, the corresponding coverage and allele frequencies were calculated for each of the 19,850,891 loci and subsequently used to filter the data. Only positions with a frequency of the major allele smaller than 0.90 and coverage larger than or equal to 350 (250 for chromosome X) were retained. 486,090 out of 19,850,891 loci (2.45%) complied with these thresholds. Supplemental Table 1.2 shows the number of SNP positions that were retained after filtering as well as the fraction per chromosome.

### Detection of monoallelically methylated loci

Likely sequencing errors in the list of filtered loci were adjusted (see Methods section 3.3 and Supplemental Methods section 2.1.3). Corrected data (available as Supplemental Data) were subsequently analysed using the developed statistical methodology. If the p-value obtained for a locus was smaller than the p-value corresponding with an FDR of 0.1 (p-value = 0.000016), the monoallelic methylation on this locus was called significant. This was true for 80 loci (see Table 1). Figure 2 depicts the genomic distribution of these 80 monoallelically methylated loci.

**Table 1.** Monoallelic DNA-methylation per chromosome. The first two columns show the specific chromosome (Chr) and the number of input entries for the statistical analysis. The third and fourth columns show the amount of loci, which obtained a p-value smaller than (or equal to) 0.005 (after first iteration) and 0.000016 (after second iteration, corresponding with FDR = 0.1), respectively.

| Chr | Input entries | $p \leq 0.005$ | $p \leq 0.000016$ |
| --- | --- | --- | --- |
| 1 | 37,259 | 98 | 3 |
| 2 | 34,184 | 113 | 10 |
| 3 | 22,065 | 73 | 4 |
| 4 | 26,442 | 68 | 3 |
| 5 | 20,500 | 54 | 1 |
| 6 | 25,708 | 112 | 3 |
| 7 | 31,450 | 153 | 8 |
| 8 | 20,868 | 63 | 0 |
| 9 | 19,908 | 78 | 0 |
| 10 | 30,138 | 108 | 7 |
| 11 | 19,255 | 82 | 8 |
| 12 | 20,484 | 51 | 0 |
| 13 | 11,709 | 88 | 3 |
| 14 | 12,297 | 40 | 1 |
| 15 | 11,829 | 47 | 2 |
| 16 | 29,432 | 64 | 4 |
| 17 | 24,824 | 158 | 3 |
| 18 | 11,783 | 29 | 1 |
| 19 | 25,066 | 96 | 8 |
| 20 | 16,201 | 45 | 7 |
| 21 | 11,977 | 40 | 2 |
| 22 | 15,012 | 70 | 2 |
| X | 7699 | 27 | 0 |
| TOTAL | 486,090 | 1757 | 80 |

**Figure 2.**
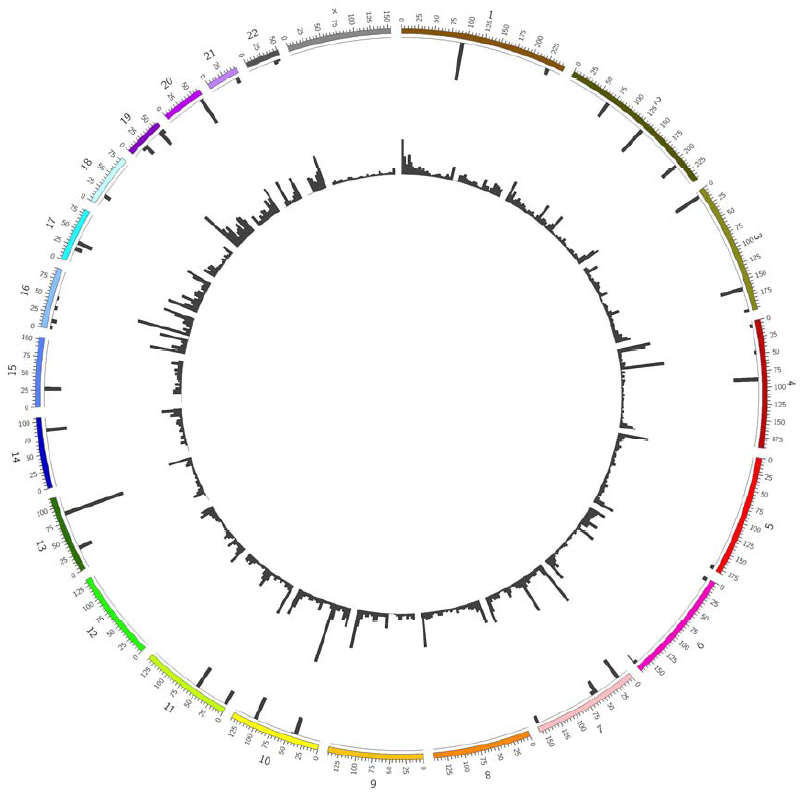
Circular representation of the genomic distribution of the 80 loci for which the monoallelic DNA-methylation was called significant. Chromosomes are divided in regions of 5,000,000 bp. The inner circle shows the histogram of all SNPs found in a specific region, whereas the outer circle shows the histograms of the significant SNPs in that same region, normalized to the number of SNPs found in that region.

In a next step, the functional location of the 80 loci with significant monoallelic DNA-methylation was determined. These results are shown in Table 2. Table 3 provides an overview of the genes in which a significant SNP position was found. Thus, of the 80 detected loci, 49 are located in a genic region (i.e. promoter, exon, intron) of which 25 are located in regions with (some) evidence, i.e. monoallelic expression, of imprinting. (Morison et al. 2001; Zhang et al. 2010, http://www.geneimprint.com).

**Table 2.** Functional annotation of the 80 loci with significant monoallelic DNA-methylation. Next to the functional annotation, the corresponding (Ensembl) gene ID is shown (GeneID).

| Chr | SNP position | Functional annotation | GeneID | Chr | SNP position | Functional annotation | GeneID |
| --- | --- | --- | --- | --- | --- | --- | --- |
| 1 | 90844629 | INTERGENIC |  | 11 | 2721568 | INTRON | ENSG00000053918 |
| 1 | 90844662 | INTERGENIC |  | 11 | 51579458 | INTERGENIC |  |
| 1 | 228757936 | PROMOTER | ENSG00000200624 | 13 | 31481030 | INTRON | ENSG00000102802 |
| 1 | 228757936 | INTRON | ENSG00000183929 | 13 | 85969909 | INTRON | ENSG00000226317 |
| 2 | 69347244 | INTRON | ENSG00000169604 | 13 | 85969941 | INTRON | ENSG00000226317 |
| 2 | 133018988 | PROMOTER | ENSG00000233786 | 14 | 88237822 | INTRON | ENSG00000258807 |
| 2 | 133020085 | EXON | ENSG00000233786 | 15 | 25123472 | INTRON | ENSG00000128739 |
| 2 | 133023691 | INTERGENIC |  | 15 | 25201659 | INTRON | ENSG00000128739 |
| 2 | 133023703 | INTERGENIC |  | 15 | 25201659 | INTRON | ENSG00000214265 |
| 2 | 133023711 | INTERGENIC |  | 16 | 3493495 | EXON | ENSG00000167981 |
| 2 | 133029769 | INTERGENIC |  | 16 | 3493495 | PROMOTER | ENSG00000122390 |
| 2 | 133032580 | INTERGENIC |  | 16 | 11415785 | INTRON | ENSG00000175643 |
| 2 | 133033524 | INTERGENIC |  | 16 | 33960762 | PROMOTER | ENSG00000207986 |
| 2 | 207122438 | INTERGENIC |  | 16 | 46411729 | INTERGENIC |  |
| 3 | 5156139 | INTERGENIC |  | 17 | 22252007 | INTERGENIC |  |
| 3 | 5156149 | INTERGENIC |  | 17 | 22259640 | INTERGENIC |  |
| 3 | 162561619 | INTERGENIC |  | 17 | 31340444 | EXON | ENSG00000108684 |
| 3 | 196722009 | INTRON | ENSG00000163975 | 18 | 18517029 | INTERGENIC |  |
| 4 | 7635629 | INTRON | ENSG00000184985 | 19 | 15279411 | INTRON | ENSG00000074181 |
| 4 | 49099668 | INTERGENIC |  | 19 | 24184564 | PROMOTER | ENSG00000251948 |
| 4 | 89618837 | INTRON | ENSG00000138641 | 19 | 54040861 | PROMOTER | ENSG00000130844 |
| 4 | 89618837 | EXON | ENSG00000177432 | 19 | 54040861 | INTRON | ENSG00000130844 |
| 5 | 178650557 | INTRON | ENSG00000087116 | 19 | 54040861 | EXON | ENSG00000130844 |
| 6 | 3849305 | PROMOTER | ENSG00000145945 | 19 | 54041242 | PROMOTER | ENSG00000130844 |
| 6 | 3849305 | INTRON | ENSG00000238158 | 19 | 54041242 | INTRON | ENSG00000130844 |
| 6 | 168784228 | INTERGENIC |  | 19 | 54057156 | INTRON | ENSG00000130844 |
| 6 | 170055316 | INTRON | ENSG00000184465 | 19 | 54057156 | PROMOTER | ENSG00000130844 |
| 7 | 19534519 | INTRON | ENSG00000223838 | 19 | 54057515 | INTRON | ENSG00000130844 |
| 7 | 56437045 | INTERGENIC |  | 19 | 54057515 | PROMOTER | ENSG00000130844 |
| 7 | 57554497 | INTERGENIC |  | 19 | 54057777 | INTRON | ENSG00000130844 |
| 7 | 61080848 | INTERGENIC |  | 19 | 54057777 | PROMOTER | ENSG00000130844 |
| 7 | 64895556 | INTERGENIC |  | 19 | 57350463 | INTRON | ENSG00000198300 |
| 7 | 157923845 | INTRON | ENSG00000155093 | 19 | 57350463 | INTRON | ENSG00000259486 |
| 7 | 158041458 | INTRON | ENSG00000155093 | 20 | 55658479 | INTERGENIC |  |
| 7 | 158041459 | INTRON | ENSG00000155093 | 20 | 55658481 | INTERGENIC |  |
| 10 | 39086215 | INTERGENIC |  | 20 | 57415110 | INTRON | ENSG00000235590 |
| 10 | 39086885 | INTERGENIC |  | 20 | 57415110 | EXON | ENSG00000235590 |
| 10 | 39086905 | INTERGENIC |  | 20 | 57415110 | PROMOTER | ENSG00000087460 |
| 10 | 39086907 | INTERGENIC |  | 20 | 57415110 | EXON | ENSG00000087460 |
| 10 | 42800026 | INTERGENIC |  | 20 | 57426449 | PROMOTER | ENSG00000235590 |
| 10 | 102279294 | INTRON | ENSG00000075826 | 20 | 57426449 | PROMOTER | ENSG00000087460 |
| 10 | 102279294 | INTRON | ENSG00000255339 | 20 | 57426449 | INTRON | ENSG00000087460 |
| 10 | 102279294 | INTRON | ENSG00000166136 | 20 | 57426726 | PROMOTER | ENSG00000235590 |
| 10 | 102279295 | INTRON | ENSG00000075826 | 20 | 57426726 | PROMOTER | ENSG00000087460 |
| 10 | 102279295 | INTRON | ENSG00000255339 | 20 | 57426726 | INTRON | ENSG00000087460 |
| 10 | 102279295 | INTRON | ENSG00000166136 | 20 | 57427132 | PROMOTER | ENSG00000235590 |
| 11 | 2019496 | PROMOTER | ENSG00000130600 | 20 | 57427132 | PROMOTER | ENSG00000087460 |
| 11 | 2019496 | INTRON | ENSG00000130600 | 20 | 57427132 | INTRON | ENSG00000087460 |
| 11 | 2019496 | PROMOTER | ENSG00000211502 | 20 | 57431165 | INTRON | ENSG00000087460 |
| 11 | 2019618 | PROMOTER | ENSG00000130600 | 20 | 57431165 | EXON | ENSG00000087460 |
| 11 | 2019618 | INTRON | ENSG00000130600 | 21 | 40757887 | INTRON | ENSG00000160183 |
| 11 | 2019618 | PROMOTER | ENSG00000211502 | 21 | 40757887 | INTRON | ENSG00000182093 |
| 11 | 2021164 | INTRON | ENSG00000130600 | 21 | 40757887 | PROMOTER | ENSG00000182093 |
| 11 | 2021206 | INTRON | ENSG00000130600 | 21 | 44011806 | INTRON | ENSG00000183486 |
| 11 | 2021980 | INTRON | ENSG00000130600 | 22 | 42078666 | PROMOTER | ENSG00000100138 |
| 11 | 2022023 | INTRON | ENSG00000130600 | 22 | 42078666 | INTRON | ENSG00000100138 |
| 11 | 2721568 | PROMOTER | ENSG00000258492 | 22 | 49077801 | INTRON | ENSG00000219438 |

**Table 3.**
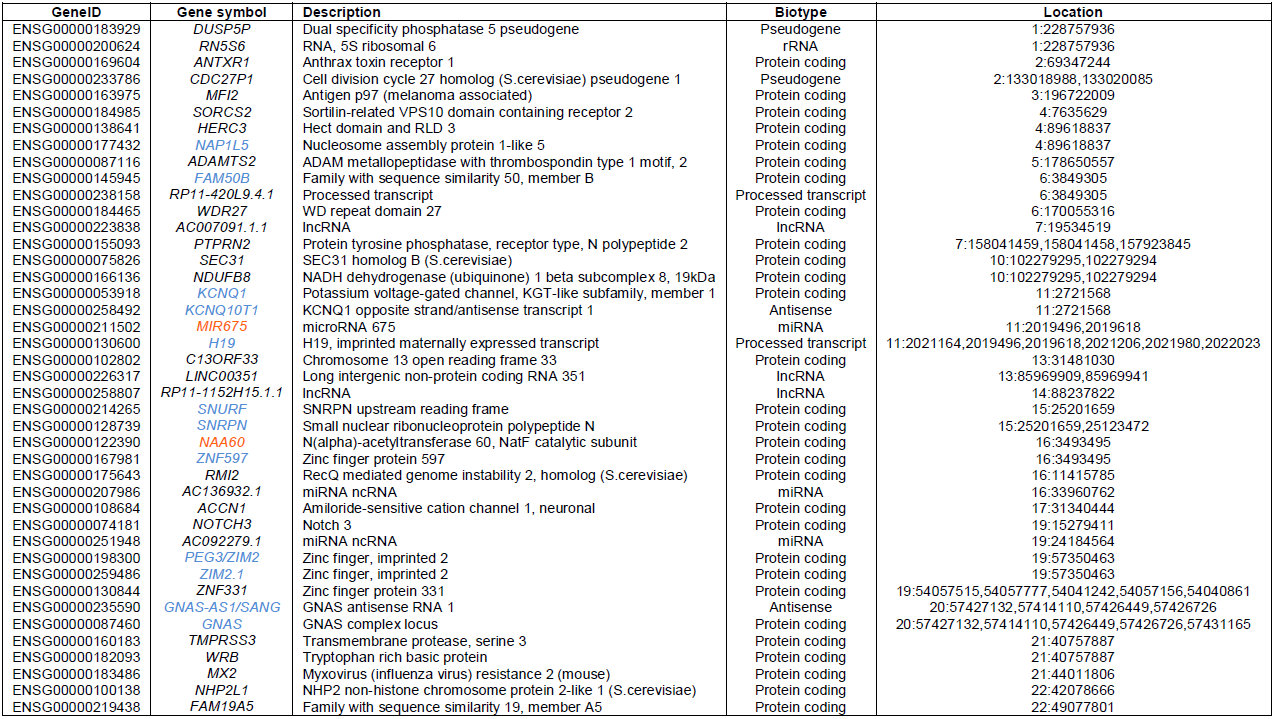
SNPs featured by monoallelic methylation located in a gene associated region with indication of Location (chromosome:location), (Ensembl) Gene ID, Gene symbol, Description and Biotype. Known imprinted genes are shown in blue, predicted imprinted genes are shown in red.

### Functional enrichment of loci with significant monoallelic DNA-methylation

Figure 3(a) represents the relative number of the different functional annotations of these 80 loci. No significant enrichment was found when genic regions were compared to intergenic regions (data not shown). The majority of the significant SNP positions are located in intronic (43.33%) and intergenic regions (37.5%). Additionally, a significant number was found in the promoter regions (13.96%). A minority of 5.21% mapped to exonic regions. In order to investigate whether one of these functional locations was under- or overrepresented compared to random data, we also performed an enrichment analysis. Figure 3(b) shows the mean classification of SNPs after 1,000 random samplings. By comparing the outcome of this random sampling with the functional locations of the 80 significant loci (see Methods section 5), the analysis indicated a significant enrichment in promoter methylation (p = 0.002), but not in other functional locations.

**Figure 3.**
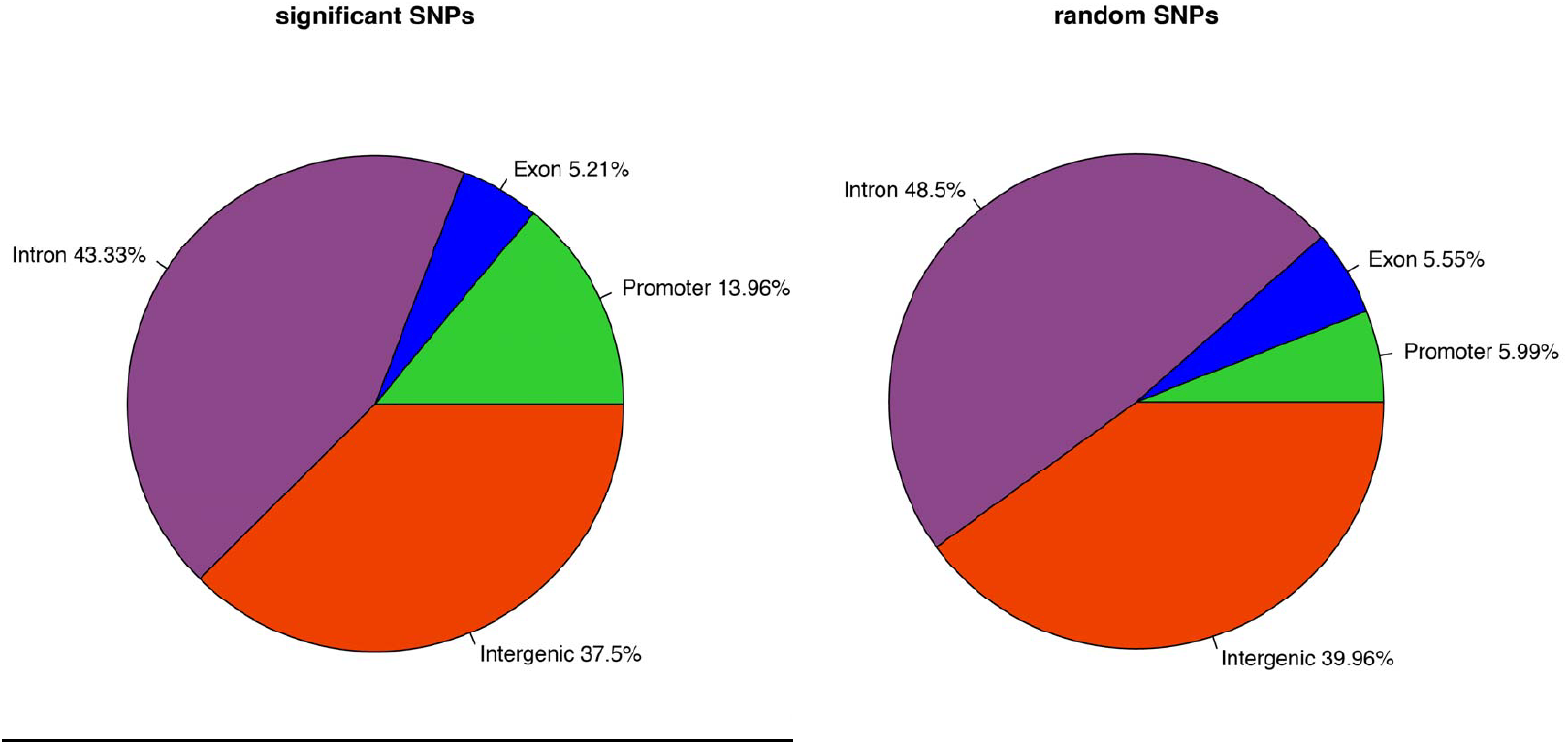
Pie charts representing the relative number of significant SNPs in the different functional classes (i.e. promoter, exon, intron and intergenic). (a) Functional classification of the significant SNPs (i.e. loci with significant monoallelic DNA-methylation). (b) Functional classification of random SNPs resulting from 1,000 iterations.

### Validation of putative loci using 14 whole-genome bisulfite sequencing data sets

After preprocessing the 14 WGBS data sets as outlined in Methods section 6, 44 out of the 80 significant were covered by at least one heterozygous sample. Table 4 summarizes both the global and the sample-specific p-values obtained for each of these 44 loci. 29 loci (65.9%) had a global p-value lower than 0.05 of which 24 (54.5%) even had a global p-value virtually equal to 0 suggesting monoallelic methylation in at least one of the 14 samples.

**Table 4.** Outcome of the additional validation of the putative loci with 14 whole-genome bisulfite sequencing (WGBS) data sets. Global and sample-specific p-values are shown for the 44 SNP loci (Chr:SNP location) that were covered by at least one heterozygous WGBS sample. Value ‘-’ in the sample columns indicates that the sample did not cover or was not heterozygous for the corresponding SNP loci. Again, known imprinted regions are shown in blue, predicted imprinted regions are shown in red.

| Chr:<br>SNP position | Global<br>p-value | Colon | Colon<br>Tumor | Cortex<br>Normal1 | Cortex<br>Normal2 | Cortex<br>AD1 | Cortex<br>AD2 | fFF | IMR90 | HepG2 | HSCP | bcell | dMesenchy | dNPC | dMEEN |
| --- | --- | --- | --- | --- | --- | --- | --- | --- | --- | --- | --- | --- | --- | --- | --- |
| 1:228757936 | 0 | 0.0195 | - | 0.387 | - | 0.2695 | 0 | 0 | 0 | 0 | 0 | 0.122 | 0 | 0 | 0 |
| 2:133018988 | 0.248 | 1 | 0.5405 | 0.525 | 1 | 1 | 0.1735 | - | - | - | - | - | 0.0685 | - | 0.482 |
| 2:133020085 | 0.0005 | 0.002 | 0.7665 | 0.7875 | 0.117 | 0.2655 | 1 | 1 | 1 | - | 0.029 | 0.124 | 0.0925 | - | 0.3705 |
| 2:207122438 | 0 | 0.1345 | 0.325 | 0.002 | 0.4405 | - | 0.0125 | 0.3495 | 0.1685 | 0.0015 | 0.5 | - | 0.03 | 0.155 | 0.0015 |
| 2:69347244 | 0.582 | - | - | 1 | - | 0.541 | - | - | - | - | - | - | - | - | - |
| 2:133033524 | 0.2535 | - | - | - | - | - | - | 0.28 | - | - | 0.6135 | - | - | - | - |
| 2:133029769 | 0.4335 | 1 | - | - | - | - | 1 | 0.175 | - | - | - | - | - | - | - |
| 2:133032580 | 0 | 0 | 0 | 0.7585 | 0.119 | 0.5695 | 0.034 | 0 | 0 | 0 | 0 | 0 | 0 | - | 0 |
| 3:162561619 | 0 | - | - | - | 0 | - | - | 0 | - | - | 0.0265 | - | - | - | 0.5525 |
| 4:7635629 | 0 | 0.0355 | - | 0.6355 | - | - | 0 | 0.004 | - | - | - | - | 0.00555 | - | 0.0005 |
| 4:89618837 | 0 | - | - | - | 0.0005 | 0 | 0.001 | 0.33 | 0 | - | - | 0.0035 | 0 | - | 0.0035 |
| 4:49099668 | 0 | 0.002 | 0.0015 | 0.034 | 0.0755 | 0.151 | 0 | 0 | 0.0595 | 0.0015 | 0.004 | 0.3295 | 0.089 | - | - |
| 5:178650557 | 0 | - | - | - | 0 | - | - | - | - | - | 0.3215 | - | 0.0045 | 0.226 | - |
| 6:170055316 | 0 | - | - | 0 | - | - | - | 0.002 | - | 0.2125 | 0.4965 | 0.566 | 0 | - | 0.2345 |
| 6:3849305 | 0 | 0 | 0 | 0 | 0.0065 | 0 | 0.0025 | 0.6925 | 0.48 | 0.0025 | - | 0 | 1 | 0.1505 | - |
| 6:168784228 | 0 | 0 | 0.04 | - | - | 0.0315 | 0 | 0.0005 | 0.0025 | 0 | - | - | 0.003 | - | - |
| 7:64895556 | 0.0265 | - | 0.38 | - | - | - | 1 | 0.0005 | - | - | - | - | - | 0.075 | - |
| 7:157923845 | 0.3995 | - | - | - | - | - | - | 0.3995 | - | - | - | - | - | - | - |
| 7:61080848 | 1 | - | - | - | - | - | - | - | - | - | 1 | 1 | - | - | - |
| 7:56437045 | 0 | - | - | 0 | - | - | - | 0 | - | - | - | - | - | 0.001 | 0.0775 |
| 7:19534519 | 0.307 | - | - | - | - | - | - | 1 | - | - | 0.3905 | - | - | - | 0.33 |
| 7:57554497 | 0.0205 | - | - | - | - | - | - | 0.0275 | - | - | 0.269 | - | - | - | 0.574 |
| 10:42800026 | 0.5405 | - | - | - | - | - | - | 1 | - | - | - | 1 | - | - | 0.27 |
| 11:2721568 | 0 | 0 | 0 | 0 | 0 | 0 | 0 | 0 | - | - | 0 | 0.1275 | - | - | - |
| 11:51579458 | 0.5285 | - | - | - | - | - | - | 0.5285 | - | - | - | - | - | - | - |
| 13:31481030 | 0.096 | 0.1905 | - | - | - | - | - | - | 0.174 | - | - | - | - | - | - |
| 14:88237822 | 0.17 | 0.184 | - | - | 0.2815 | 1 | - | 0.1725 | - | - | - | 0.0445 | - | - | 0.6605 |
| 15:25123472 | 0 | 0 | 0 | 0.0475 | 0.003 | 0 | 0 | 0.4215 | 0.314 | - | 0.0045 | 0 | - | - | - |
| 15:25201659 | 0 | - | - | 0 | 0 | - | - | 0.5625 | - | - | 0.4805 | 0.012 | 0 | 0 | 0 |
| 16:46411729 | 0 | 1 | 0.6475 | - | - | - | - | 0 | - | 0.546 | - | - | - | - | - |
| 16:3493495 | 0 | - | - | 0 | 0 | - | 0.0355 | 0 | 0 | - | 0.861 | 0.0095 | 0 | - | 0.001 |
| 16:11415785 | 0.002 | - | - | - | - | - | - | 0.0475 | 0 | - | - | - | - | - | - |
| 17:22252007 | 0.74 | 1 | - | 0.6095 | - | - | 0.627 | - | - | - | 0.6155 | - | - | - | - |
| 17:22259640 | 0.7215 | 0.8215 | 0.7795 | 0.3425 | 1 | 1 | 0.613 | 0.3785 | 0.6265 | - | 0.212 | 0.7165 | 1 | - | - |
| 18:18517029 | 0 | - | 0.0415 | - | 0.017 | 0.1795 | 0.0045 | 0 | - | - | 0 | 0.307 | - | - | 0 |
| 19:15279411 | 0 | - | - | 0.299 | 0.054 | 0 | 0.0005 | 0 | - | 0.568 | 0.839 | 0.0155 | 0 | - | 0.016 |
| 19:57350463 | 0 | 0 | 0.0005 | 0 | 0 | 0 | - | 0.004 | - | 0 | 0.2945 | 0.0095 | 0 | 0 | 0 |
| 19:24184564 | 0 | - | - | - | - | - | - | 1 | - | - | - | - | - | 0 | 0.0005 |
| 20:57415110 | 0 | - | - | 0.016 | 0 | - | 0 | 0 | 0.0095 | 0.4675 | - | 0 | 0 | - | 0 |
| 20:57431165 | 0.0625 | - | - | 0.5545 | - | - | - | 0.011 | 0.8245 | - | - | - | - | - | - |
| 21:44011806 | 0.0145 | - | - | - | - | - | - | 0.0145 | - | - | - | - | - | - | - |
| 21:40757887 | 0 | 0 | 0 | - | - | - | - | 0.0255 | - | - | 0 | - | 1 | - | - |
| 22:42078666 | 0 | 0 | 0.005 | - | 0 | - | - | - | - | - | - | - | 0 | - | 0.013 |
| 22:49077801 | 0.715 | - | - | - | - | - | - | 0.715 | - | - | - | - | - | - | - |

## DISCUSSION

Monoallelic gene expression is typically initiated early in the development of an organism and stably maintained. Erroneous monoallelic expression has been related to several non-Mendelian inherited genetic disorders. DNA-methylation plays a significant role in the regulation of monoallelic expression. The choice of which allele will be monoallelically expressed can be either random or a priori defined by imprinting. Here we introduced a methodology to screen for genes that exhibit monoallelic DNA-methylation and thus might regulate monoallelic expression.

Using MethylCap-seq, methylome profiles of 334 samples, mostly human cancer samples of diverse origin, were obtained. In summary, upon extra filtering and data correction, for each SNP locus the Hardy-Weinberg theorem was applied to evaluate whether the observed frequency of samples featured by biallelic methylation is lower than randomly expected. Using a permutation approach, loci with a p-value smaller than the p-value corresponding with a selected FDR of 0.1 were assumed to be monoallelically methylated. Finally, this resulted in the identification of 80 loci that showed significant monoallelic DNA-methylation.

Functional location of these monoallelic events might provide deeper insight in the unraveling of monoallelic mechanisms and are provided in Tables 2 and 3. It is common that imprinted genes are present within clusters and share common regulatory elements, such as non-coding RNAs and differentially methylated regions (DMRs). If these DMRs control the imprinting of one or more genes, these regions are called imprinting control regions (ICRs). It is known that many of these ICRs are located in intergenic regions. As some of the found loci are located in intergenic regions as well as in known (long) non-coding RNAs (lncRNAs) (see Tables 2 and 3, respectively), it is possible that these regions present new regulatory elements involved in imprinting. Furthermore, when we take a closer look at the intergenic regions, the SNP on chromosome 2 with position 207,122,438 also shows significant monoallelic methylation. This is interesting, because this locus falls in *GPR1AS*, a recently found imprinted lncRNA in the *GPR1-ZDBF2* intergenic region (Kobayashi et al. 2013), corroborating the outcome of this study and indicating that the intergenic regions are also of interest for further analyses. Of the 80 loci, 49 were located in genic regions of which 25 are already linked to imprinting, further strengthening our results. For example, on chromosome 11 (2.01-2.03 Mb), the *IGF2/H19* region was highlighted with 6 SNPs (Figure 2). This locus is a well known imprinted region that’s also linked to the Beckwith-Wiedemann syndrome and Wilms’ tumor (Steenman et al. 1994; Egger et al. 2004; Robertson 2005; Hubertus et al. 2011). The *H19* gene codes for a lncRNA of which expression is negatively correlated with the expression of the neighbouring gene insulin-like growth factor 2 (*IGF2*). Usually the paternal copy of *H19* is methylated and silent, while the maternal copy is hypo- or unmethylated and expressed. The same is true for the imprinted region on chromosome 15 that is correlated to the Prader-Willi syndrome (20.7-30.3 Mb) (Egger et al. 2004). In *SNRPN*, one of the genes in this region where loss of imprinting is linked to the Prader-Willi syndrome, 2 significant SNPs were identified. For a couple of genes (or regions), like for example *H19*, more than one significant SNP locus was found. Because some of these SNPs are in a distance of more than 400 bp (the cut-off length of sequence reads during mapping) of each other, these prove independently the presence of monoallelic DNA-methylation in that particular region. These SNPs thus provide ‘multiple proof’ in the identification of the particular monoallelically methylated region and lend added value to the results. Not unexpectedly, Figure 3 and the enrichment analysis clearly demonstrated enrichment for monoallelic methylation in promoter regions, although it should be noted that this enrichment is rather limited in absolute number.

Further validation was performed using 14 publicly available WGBS data sets. Of the 80 significant loci, 44 were covered by a heterozygous sample and could thus be further examined. 29 of the 44 loci (65.9%) obtained a global p-value lower than 0.05 of which 24 (54.5%) a global p-value of virtually 0, indicating monoallelic methylation in one or more samples. For only 9 of these 24 SNP loci evidence of imprinting already exists, so that with this subset of 14 WGBS samples at least 15 new monoallelically methylated regions, found with our new data-analytical framework, are validated.

There are a couple of important remarks that come with the proposed methodology: i) The basic assumption that MethylCap-seq data from biallelically methylated loci are generally in Hardy-Weinberg equilibrium only holds for samples originating from a panmictic population (i.e. a single population that is long-term randomly mating). If this is not the case and the samples are not panmictic, this could possibly give rise to some false positives. Thus, for samples that slightly deviate from the assumption of panmixia, an extra validation of the resulting loci is necessary to assure qualitative results (as was done in this study). ii) The approach doesn’t take into account that loci with monoallelic methylation will be picked up less efficiently than biallelic loci resulting in less power leading to a less efficient detection of monoallelically methylated loci. By consequence, the methodology is less sensitive and thus too conservative, though this has no effect on the reliability of those results deemed significant. iii) To eliminate sequencing errors as well as to reduce the computational time and effort a filtering step was performed. Consequently, some data will not be analysed and this could interfere with the detection of monoallelic DNA-methylation. However, the benefits of filtering outweigh the possible drawbacks: the computational load reduces significantly and loci that do not pass the filter cut-off will have insufficient power to be detected. iv) The approach used to correct for possible sequencing errors disfavors the presence of monoallelic DNA-methylation: only two correction rounds were performed and the sequencing error estimate of 0.25% is the lowest estimate reported (Quail et al. 2012). But although the correction method can be considered a bit too stringent, it will assure a better quality of the obtained results and will not give rise to more false positives. In fact, it will possibly reduce the amount of false negatives and thus allows a more sensitive identification of monoallelically methylated loci that would otherwise not have been detected. v) The analysis was performed with samples originating from different tissues that were mostly cancer tumors. The fact that tumors are epigenetically less stable (Varley et al. 2013), makes it probably more difficult to detect monoallelic methylation.

Although we opted to use a stringent approach, the outcome clearly demonstrates that our methodology is still sensitive enough and produces satisfying results of high quality. Because the methodology used a mixture of samples originating from different tissues and demanded a high overall coverage, the identified loci are presumably monoallelically methylated in a variety of cell types, underlining their importance and biological relevance. In conclusion, the obtained results prove that the proposed methodology is effective. In the future, it would also be very informative to repeat the analysis on samples that are of normal origin and in extent from the same tissue. The latter would be very valuable in the study of tissue-specific monoallelic DNA-methylation, expression and thus tissue-specific cell differentiation. Next to MethylCap-seq, our approach also opens the door to other applications, like ChIP-seq-based detection of monoallelic protein-DNA binding events and histone modifications.

## DATA ACCESS

The filtered and corrected SNP data used as starting data for our developed methodology is made available as Supplemental Data.

## ACKNOWLEDGMENTS

We would like to thank the N2N “nucleotides 2 networks” Multidisciplinary Research Partnership) for financial support. S. Steyaert also wants to acknowledge the IUAP for funding.

